# Caller identification and characterization of individual humpback whale acoustic behavior

**DOI:** 10.1101/2023.10.17.562793

**Authors:** Julia M Zeh, Valeria Perez-Marrufo, Dana L Adcock, Frants H Jensen, Kaitlyn J Knapp, Jooke Robbins, Jennifer E Tackaberry, Mason Weinrich, Ari S Friedlaender, David N Wiley, Susan E Parks

## Abstract

Acoustic recording tags are biologging tools that provide fine scale data linking acoustic signaling with individual behavior; however, when an animal is in a group, it is challenging to tease apart calls of other conspecifics and identify which individuals produce each call. This, in turn, prohibits robust assessment of individual acoustic behavior including call rates and silent periods, call bout production within and between individuals, and caller location. To overcome this challenge, we simultaneously instrumented small groups of humpback whales on a western North Atlantic feeding ground with sound and movement recording tags. This simultaneous tagging approach enabled us to compare the relative amplitude of each call across individuals and infer caller identity though amplitude differences. Focusing on periods when the tagged animals were isolated from other conspecifics, we were able to assign caller ID for 97% of calls in this dataset. From these labeled calls, we found that humpback whale individual call rates are highly variable across individuals and groups (0-89 calls/h), with calls produced throughout the water column and in bouts with short inter-call intervals (ICI = 2.2 s). Most calls received a likely response from a conspecific within 100 s. These results are important for modelling signal detection range for passive acoustic monitoring and density estimation. Future studies can expand on these methods for caller identification and further investigate the nature of sequence production and counter-calling in humpback whale social calls. Finally, this approach can be helpful for understanding intra-group communication in social groups across other taxa.

**Summary statement:** Tagging entire humpback whale social groups with sound and movement recording tags allows us to for the first time parse out call behavior within groups and understand individual acoustic behavior.

## Introduction

In studies of animal communication, it is valuable to be able to differentiate the sender and receiver of a given signal (Demartsev et al. 2022). Once caller identity has been assigned, more detailed information about the vocal behavior of a species can be inferred, including individual call rates, timing of signal production, and the production of acoustic sequences within and between individuals. However, in naturalistic social settings across taxa in both the lab and in the field, assigning acoustic signals to the individual that produced them can be challenging (Heckman et al. 2017, Stimpert et al. 2020). Unless an animal gives an obvious visual cue when vocalizing, caller identification requires either highly precise sound source localization (e.g., Miller et al. 2004, Heckman et al. 2017) or some other method of differentiating the calls of one individual from those of conspecifics in the vicinity. Animal-borne tags containing movement and acoustic sensors provide valuable fine-scale data to link individual sound production and behavior (Johnson et al. 2009). However, these acoustic sensors record all detectable sounds from both the tagged animal and nearby conspecifics (Johnson et al. 2009). In social groups, conspecifics are often in close proximity to the tagged animal; therefore, calls from other animals present challenges for caller identification. This is especially problematic for studies of social animals and of underwater sound production, since sound propagates efficiently and rapidly through water, resulting in a high probability of detecting nearby vocalizing conspecifics.

While past studies have used various methods for caller identification, most of these methods remain problematic or are limited to only certain taxa. For example, the angle of arrival of recorded sounds on stereo hydrophones in tags have been used for caller ID (Johnson et al. 2006, Madsen et al. 2013, Oliveira et al. 2013, Kragh et al. 2019), sometimes in concert with separations from the social group (Jensen et al. 2011, Perez et al. 2017). While calculations of the angle of arrival of sounds have been useful for assigning calls as focal (i.e., from the tagged animal) or non-focal for the high frequency clicks and whistles of odontocete species (Johnson et al. 2006, Madsen et al. 2013, Oliveira et al. 2013, Kragh et al. 2019), these methods prove problematic for low frequency baleen whale calls, whose longer wavelengths and gradual amplitude onset hinder localization with narrow inter-hydrophone spacing.

The use of individual identity information in the recorded sounds is possible for caller ID for some species. For example, individual spectral features in goat (*Capra aegagrus hircus)* vocalizations have allowed for caller identification (O’Bryan et al. 2019), as has the inter-pulse interval in sperm whale (*Physeter macrocephalus)* codas, which can be used to infer body size (Schulz et al. 2011, Gero et al. 2016). These methods are only possible in select situations when animal vocalizations contain individual identity cues, and these cues are known. No such methods currently exist for robust individual identification from baleen whale calls.

In contrast, signal-to-noise ratio (SNR) thresholds have been used frequently in assigning caller ID in baleen whale tag data (e.g., Oleson et al. 2007, Parks et al. 2011), but this method can be problematic given that individuals vary the source level of their sounds; quiet sounds may come from the tagged animal and nearby conspecifics may produce calls detected on the tag with high SNR (Stimpert et al. 2020). SNR measurements will also depend on tag attachment location as well as on call type and background masking noise.

Finally, some studies have used signatures of very low-frequency sounds picked up by the tag accelerometer data for caller ID (Goldbogen et al. 2014, Stimpert et al 2015, Saddler et al. 2017, Stimpert et al 2020). While accelerometer signatures of calling behavior have shown promise, these methods can still be ambiguous since accelerometers have been shown to pick up calls from both the tagged whale and from nearby conspecifics (Saddler et al. 2017). Furthermore, sufficiently high-resolution accelerometer data would be necessary to detect higher frequency baleen whale calls and even then, the mechanism involved in accelerometer detection of vocalizations is still unclear and not all focal calls register on the accelerometer (Stimpert et al. 2020).

More recently, Kragh et al. (2019) distinguished bottlenose dolphin (*Tursiops truncatus*) whistles produced by the tagged individual from those produced by non-focal animals via a combination of the angle of arrival of whistles and, when pairs were tightly associated, differences in call intensity recorded across the two tags. Comparisons of call amplitude across tags requires tags deployed on all individuals in a social group but shows promise for studies of baleen whale calls. Here we show how this method can be used to distinguish focal and non-focal calls in tag data from humpback whales (*Megaptera novaeangliae)*.

Humpback whales are found across the globe and migrate annually between low latitude breeding grounds and high latitude feeding grounds (Dawbin 1966). They are acoustically active throughout their range, producing a variety of social sounds across various contexts (Dunlop et al. 2007). On their feeding grounds, humpback whales can be found in large aggregations and are vocally active across different contexts (Stimpert et al. 2011). Males also produce a complex, hierarchically structured song, which is recorded most often on the breeding grounds (Payne and McVay 1971). Singing behavior is known to rely on rhythmically produced acoustic sequences and this can facilitate tracking individual singers and teasing apart individual songs (e.g., Stanistreet et al. 2013). In addition to songs, there is ample evidence of social calls produced in bouts by individuals (e.g., Rekdahl et al. 2015). In the South Pacific, migrating humpback whales were shown to produce most of their social calls in bouts with 3.9 seconds or less between calls, based on an SNR threshold for estimating which calls were focal (Rekdahl et al. 2015, Cusano et al. 2022). Bouts are widely variable in duration, context, and call types, but there is some evidence of syntactical rules governing the order of call types in a bout (Rekdahl et al. 2015).

Humpback whales are challenging subjects for caller identification because, in addition to being baleen whales with far-reaching low frequency calls, they are often vocally active in social settings, when many individuals are vocalizing near one another. Thus, little data exists that has allowed for quantitative analysis of the nature of individual bout production or of vocal exchanges. In addition to call bouts from a single individual, inter-individual call bouts are involved in vocal exchanges. While some animals exhibit simple call and response dynamics, others have shown evidence of temporal rules in call exchanges indicative of turn-taking and temporal coordination (e.g., Takahashi et al. 2013, Demartsev et al. 2018). These turn-taking rules involve limited or no interruptions and describe the periodicity of vocal exchanges, in line with similar analysis of coordination in human conversation (Takahashi et al. 2013). Group vocal coordination may also arise from individual rules related to call inhibition and excitation in response to conspecific vocalizations (Demartsev et al. 2018). In part due to challenges with caller identification, quantitative descriptions of vocal exchanges, also sometimes referred to as counter-calling, are lacking for humpback whales.

Without robust caller ID methods, it is difficult to study individual vocal behavior and calculate individual call rates. Call rates are increasingly important for passive acoustic monitoring (PAM) and acoustic density estimation (i.e., Marques et al. 2013), especially in the context of vocal exchanges. The behavioral context of signal production on an individual level, such as the depth at which animals are vocalizing, is similarly challenging to describe, but important for modelling signal detection range for use with PAM and density estimation.

In this study, we test whether we can use calls’ received levels from acoustic recording tags simultaneously deployed on all animals in a social group to assign caller identity. We then describe individual vocal behavior and explore vocal exchanges in groups (pairs and trios) of North Atlantic humpback whales on the Gulf of Maine feeding ground. Specifically, we look at how individual vocal behavior relates to individual movement behavior by calculating the depth at which individuals vocalize. Furthermore, we investigate the acoustic context of individual calls by testing for and characterizing bout production and call timing in vocal exchanges, all of which could not be assessed without robust caller identification methods.

## Materials and methods

### Data Collection

Sound and movement data were collected from humpback whales in the Gulf of Maine in and around Stellwagen Bank National Marine Sanctuary in the Western North Atlantic between 41.5 and 43.2°N and 69.3 and 70.5°W. Archival digital acoustic recording tags (Dtag version 2; Johnson and Tyack 2003) were attached via suction cups from a handheld 7-15m pole in July 2006-2009 (Wiley et al. 2011). Dtag hydrophones recorded at a sampling rate of either 64 or 96 kHz and orientation sensors recorded at a sampling rate of 50 Hz, which were decimated to 5 Hz for analysis. We did not examine the accelerometer data for signatures of vocalizations because the sampling rate of the accelerometers used in this study was too low; a 50 Hz sampling rate would only allow detection of sounds up to 25 Hz, and most humpback whale vocalizations are >100 Hz (Stimpert et al. 2011). Behavioral observations, including social affiliations, were also collected concurrently from a small inflatable vessel at a distance of a few hundred meters away (e.g., Weinrich 1991, Weinrich et al. 1992). A handheld GPS onboard the vessel was used to record the location of tag deployments. Individual whales were identified based on the unique shape and pigmentation pattern of their ventral flukes (Katona and Whitehead 1981). They were photographed and matched to photo-identification catalogues from long-term studies led by the Center for Coastal Studies and the former Whale Center of New England. Whales were classified as male or female based on molecular sex determination (Palsbøll et al. 1992, Bérubé & Palsbøll 1996), a photograph of the genital slit, or, in the case of females, a calving history (Glockner, 1983). Calves were classified based on their size, stereotypical behaviours and close, consistent association with a mature female (the mother). The age class of other individuals was assigned from longitudinal data on the exact or minimum age of each individual. With the exception of the calves, all of the individuals in the study were at least five years old and therefore considered adults (Chittleborough 1959, Clapham 1992, Robbins 2007).

### Acoustic Analysis

#### Focal Call Assignment

To ensure that we could accurately assign calls to specific individuals in the group, we only used tag data from periods of time when 1) all whales in a group were equipped with tags; 2) no untagged whales were associated or in close proximity to the group (<500m); and 3) visual observers were recording behavioral focal follow data to confirm the social associations and behavioral context of the tagged whales. Most data analysis began at the time point when the last tag in the group was deployed. Analysis ended when behavioral observations stopped, another whale joined the group, or a tag detached from one of the whales in the group. During these analysis periods, we manually detected all humpback whale vocalizations and compared the relative received level of the signal across all the tags in the group to identify which animal was calling. A call should have the greatest received level on the tag attached to the whale producing the sound, regardless of the sound source level, because that tag would be closest to the sound source.

Experienced analysts manually selected individual humpback whale calls in the acoustic record of each tag. Tag acoustic records were analyzed both individually using Raven Pro v2.0 (K. Lisa Yang Center for Conservation Bioacoustics at the Cornell Lab of Ornithology 2023) and simultaneously in MATLAB 2019b (The MathWorks Inc. 2019) using custom scripts. All humpback whale calls were selected in Raven Pro, regardless of whether the analyst thought the calls were from the tagged individual. Single and simultaneous tag audits were conducted by separate analysts and analysts were blind to the results of the analysis with the other method. All sound files were thus browsed by at least two experienced analysts to reduce false positives and false negatives. Once detections from the two analysts were combined, simultaneous tag analysis was used to identify focal (i.e., originating from the tagged whale) and non-focal (i.e., originating from a whale other than the tagged whale) calls in the tag record based on relative call intensity across tags (Kragh et al. 2019). This involved plotting spectrograms and relative intensity plots from time-aligned acoustic data from all concurrent tags (Jensen et al. in prep). For each manually selected call, the spectrogram(s) of the other tag(s) were examined for instances of the same call (Figure 1). If calls were not recorded on the other tag(s) in the group, they were assumed to be focal calls. If calls were recorded on the other tag(s) in the group, relative intensity was compared and calls were assigned as either focal (when relative intensity was highest on that tag), non-focal (when relative intensity was lower than it was on another tag), or indeterminate (when there was no clear difference in relative intensity across tags). When one tag was obscured due to noise, including surfacing noise, the call was marked focal for the tag where it was visible on the spectrogram. Indeterminate calls may have been produced by a tagged whale when in very close proximity to another tagged whale or may have been produced by a whale outside the group and recorded with the same intensity on all tags. We also noted whether calls were detected on multiple tags and whether noise (e.g., flow noise, splashing noise during a surfacing) was present on one of the tags which may have masked detection of a call.

**Figure 1:**
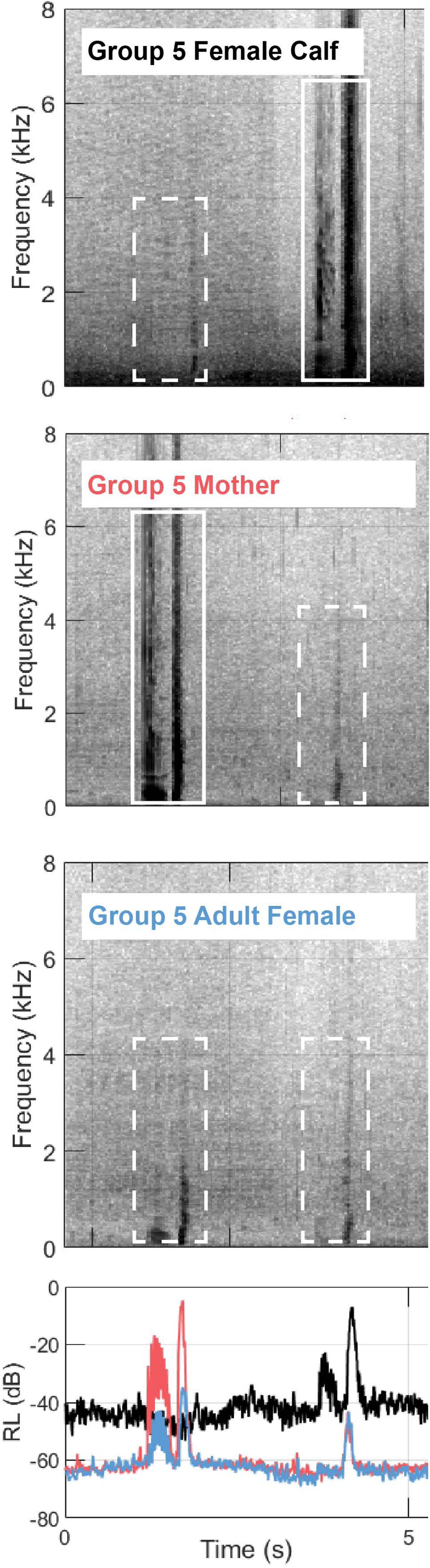
Spectrograms and received level (RL) plot showing two vocalizations recorded on all three tags in Group 5. Dashed boxes show the non-focal instances of the calls and solid boxes show the focal instance of each call. The color of the text labels on the spectrogram correspond to the colors of the lines in the RL plot.

We measured the received level (RL) of focal and non-focal calls in MATLAB by first decimating the audio to 12kHz and then applying a 500 Hz high-pass filter to reduce flow noise. We only measured received level for those focal and non-focal calls that did not overlap temporally with other sources of noise. For those calls where we could measure the signal, we measured the root-mean-squared (rms) RL using the *rms* function in MATLAB based on a 90% energy window. We then converted this value to dB re 1 µPa using a nominal hydrophone sensitivity of −171 dB re 1 V/µPa (Stimpert et al. 2011). After making RL measurements, we paired up focal and non-focal instances of the same call to measure the difference in RL of the same call when it was recorded across multiple tags. We also calculated RL differences across tags when a call was recorded on more than one tag; however, it is important to note that call RLs also depend on tag placement on an animal and variation in tag placement across deployments would thus affect these calculated differences. All statistical analyses were done in R version 4.1.2 (R Core Team 2021).

Only calls labeled focal were retained for further analysis, and we used these data as well as the analysis duration to calculate raw call rates at both the individual and group levels. We also calculated the proportion of the total analysis period that was silent (i.e., contained no call detections from any individual in the group).

#### Vocal Exchanges and Bout Analysis

To understand the communicative context of calling behavior, we investigated call timing both within and between individuals by looking at individual call bouts and inter-individual vocal exchanges. We conducted a bout analysis by calculating a bout end criterion (BEC), which determines a threshold for defining calls as part of a bout (Sibly et al. 1990). First, we calculated inter-call intervals (ICIs) from the start of one call to the start of the next call from the same individual. We then log-transformed the inter-call interval data and used the R package *diveMove* to determine the BEC using the maximum likelihood estimation method (Luque and Guinet, 2007; Luque 2007). The package *diveMove* was developed to look at dive bouts using dive intervals, but the methods are applicable for intervals and bouts of any behavioral parameters. The BEC method assumes that the distribution of behavioral data combines two or more Poisson processes, including fast processes (calls within a bout) and slow processes (calls in separate bouts). The BEC is calculated as the point where the distribution switches between these two processes and has been described as a “broken-stick” model (Sibly et al. 1990). After calculating the BEC, we classified calls with ICIs less than the BEC as bouts.

We examined vocal exchanges in groups by looking at relative call timing between individuals. We calculated the between-individual inter-call interval as the difference between the start time of a call and the start time of the next call made by a different individual. We then used these inter-individual ICI data to calculate a probability density function and integrated over the function to get the area under the curve (AUC).

#### Movement analysis

Accelerometer, magnetometer, and pressure sensor data were calibrated and processed using custom-written MATLAB scripts (animaltags.org). Depth of call production was also calculated for all focal calls across all individuals by comparing time of call production to the pressure time series from the tag. Maximum dive depths were calculated for each dive and each individual in order to investigate call production depth relative to dive depth. To assess dive and call production depth relative to bathymetry, we also report estimated seafloor depth based on GPS coordinates from where the tag was deployed on the whale.

**Table 1:** Summary of tag data, class of individuals tagged, analysis duration, and total focal calls detected. . Totals represent individual detections and are not the same as unique sounds; there is overlap in calls that are counted as focal on one tag and non-focal on another, or as indeterminate on multiple tags. For example, although there were 58 detections of indeterminate calls, this represents only 27 unique calls that could not be attributed to a specific individual.

| Date | Group | Analysis duration (hh:mm) | Whale class | Total number of focal calls | Total number of non-focal calls | Total number of indeterminate calls | Focal call rate (calls/hour) |
| --- | --- | --- | --- | --- | --- | --- | --- |
| July 19, 2006 | 1 | 1:28 | Adult Female 1 | 0 | 15 | 0 | 0 |
|  |  |  | Adult Female 2 | 20 | 0 | 0 | 13.6 |
| July 17, 2007 | 2 | 2:40 | Male Calf | 3 | 0 | 0 | 1.1 |
|  |  |  | Mother | 0 | 1 | 0 | 0 |
| July 7, 2008 | 3 | 2:27 | Adult Male | 8 | 5 | 0 | 3.3 |
|  |  |  | Adult Female | 12 | 8 | 0 | 4.9 |
| July 14, 2008 | 4 | 0:31 | Adult Female | 46 | 9 | 1 | 89 |
|  |  |  | Adult Male | 15 | 15 | 1 | 29 |
| July 22, 2009 | 5 | 3:47 | Female Calf | 335 | 173 | 10 | 88.5 |
|  |  |  | Mother | 314 | 75 | 12 | 83 |
|  |  |  | Adult Female | 139 | 113 | 4 | 36.7 |
| July 29, 2009 | 6 | 0:17 | Adult Female | 0 | 0 | 0 | 0 |
|  |  |  | Adult Male | 6 | 0 | 0 | 21.2 |
| July 20, 2009 | 7 | 6:55 | Adult Female | 15 | 28 | 11 | 2.2 |
|  |  |  | Female Calf | 18 | 37 | 11 | 2.6 |
|  |  |  | Mother | 77 | 11 | 8 | 11.1 |

## Results

In total, we analyzed 46 hours 52 minutes of tag data for which we had synchronous tags on all whales in a given group with concurrent behavioral observations, which allowed for received level comparisons and caller ID. This included 16 tags from 7 distinct groups of whales, with 12 females and 4 males. These 16 whales also included three calves and three mothers. Most of the whales were foraging for most of the tag duration, although some were also traveling or resting.

### Focal call assignment

We were able to use received level comparisons across tags (i.e., Figure 1) to identify 1008 total focal calls in the dataset. Some individuals did not produce any calls, while others called over 300 times (Table 1). We also identified 490 non-focal calls in total, which were the quietest instance of a call when it was detectable on multiple tag records. Finally, there were 27 calls (2.6%) that could not be assigned to an individual because of the similarity in received level across tags. Of the 1035 total unique calls detected across all individuals, 393 calls (38%) were detected across multiple tags, 621 calls (60%) were only detected on one tag, and there were 621 instances, a total of 489 calls (47%), when noise was present, so it was possible that a call could have been detected on multiple tags but was masked by noise. There is a chance that some calls were misclassified when noise was present because the highest RL version of the call was masked by noise and thus a lower RL non-focal call would have been marked as focal. However, the amplitude of the noise in these cases was generally low and would likely have only masked non-focal calls or some low amplitude focal calls, reducing the risk of this type of error. The average RL of all focal calls was 129 dB re 1 µPa and the average RL of all non-focal calls was 122 dB re 1 µPa. The mean difference in RL of a call recorded across tags was 15 dB. The distribution of RLs of non-focal calls overlaps entirely with the distribution of the RLs of focal calls (Figure 2).

**Figure 2:**
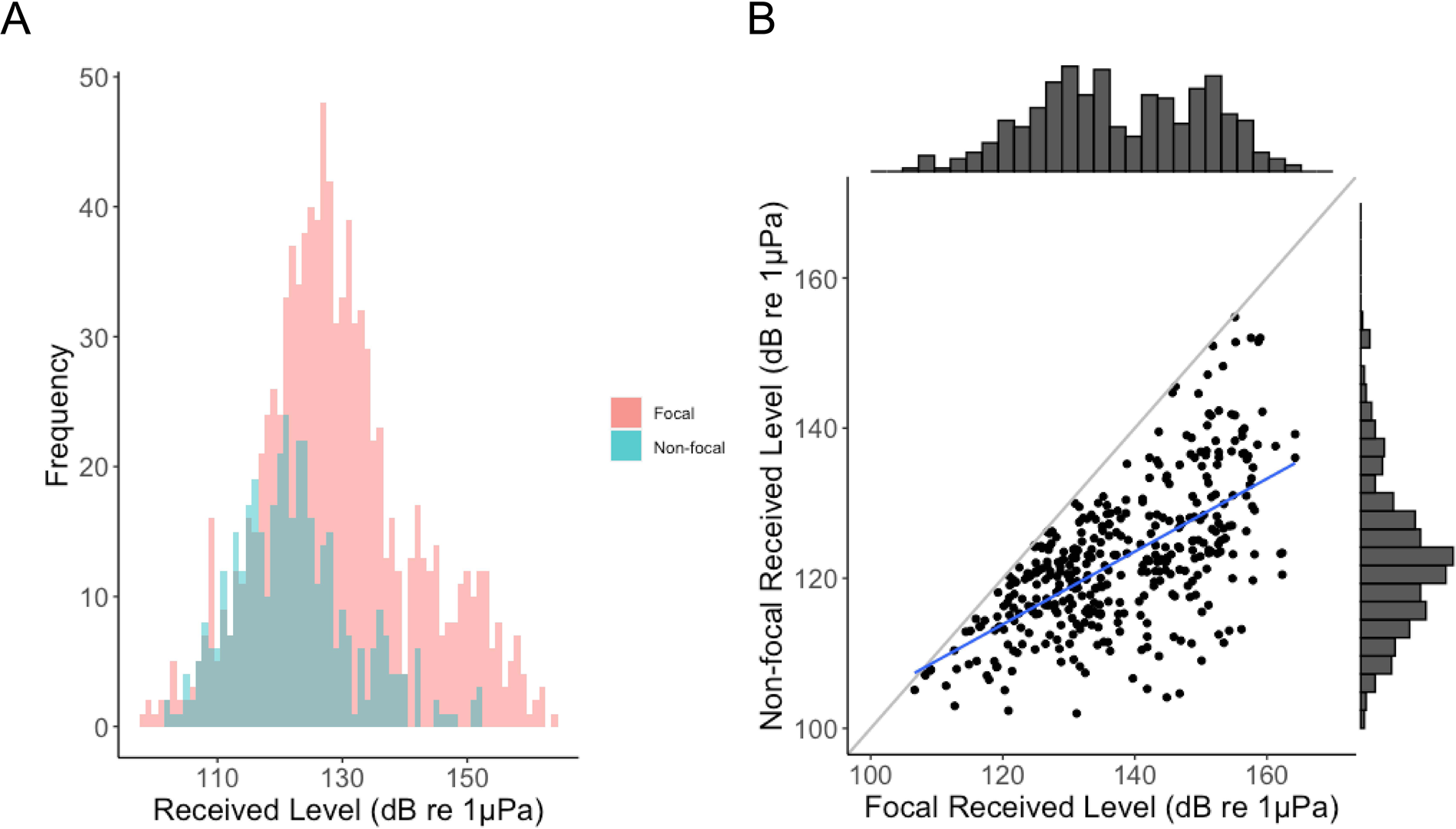
Received levels of focal and non-focal calls. A) Histogram showing the distribution of received levels of focal (red) and non-focal (teal) calls overlaid on the same plot. B) Scatterplot for calls that were recorded across multiple tags, the non-focal received level is plotted against the corresponding focal received level of the same call. The identity line is shown in gray and a linear regression line for the data is shown in blue. Marginal histograms show the distribution of focal RLs (x-axis) and non-focal RLs (y-axis).

Hourly call rate, based on the analysis duration and number of focal calls detected, ranged from 0 to 87 calls per hour (Table 1). The average call rate across individuals was 23 calls per hour and across groups was 55 calls per hour. On average across tags, 71% of the analysis period was silent and contained no call detections. The longest periods of silence across tags ranged from 278 seconds to 3.62 hours.

### Bout analysis

The BEC for this dataset is 2.2 seconds, meaning that any calls with an ICI of less than 2.2 seconds were classified as part of bouts, while those with greater ICIs were not. On average, across individuals, 79% (+/-15% SD) of calls were produced as part of bouts. Bouts were made up of 2 to 6 calls on average, and individuals produced between 0 and 69 total bouts. Bout rates ranged from about 0 to 14 bouts per hour. Inter-individual ICIs ranged from 0.05 to about 8000 seconds. The AUC between 0 and 100 seconds for the probability density function was 0.58, meaning that 58% of the time, a call from one whale was followed by a call from a different whale within 100 seconds (Figure 3).

**Table 2:** Number of bouts, bout rate, and mean number of calls per bout for all tags.

| Group | Analysis duration (hh:mm) | Whale class | Total number of bouts | Bout rate (bouts per hour) | Mean number of calls per bout |
| --- | --- | --- | --- | --- | --- |
| 1 | 1:28 | Adult Female 1 | 0 | 0 | 0 |
|  |  | Adult Female 2 | 3 | 2 | 6 |
| 2 | 2:40 | Male calf | 1 | 0.4 | 2 |
|  |  | Mother | 0 | 0 | 0 |
| 3 | 2:27 | Adult male | 2 | 0.8 | 3.5 |
|  |  | Adult female | 3 | 1.2 | 3.7 |
| 4 | 0:31 | Adult female | 7 | 13.5 | 6 |
|  |  | Adult male | 2 | 3.9 | 3.5 |
| 5 | 3:47 | Female calf | 4 | 1 | 2.8 |
|  |  | Mother | 4 | 1 | 2 |
|  |  | Adult female | 17 | 4.5 | 3.9 |
| 6 | 0:17 | Adult female | 0 | 0 | 0 |
|  |  | Adult male | 1 | 11.3 | 3 |
| 7 | 6:55 | Adult female | 69 | 6.6 | 3 |
|  |  | Female calf | 40 | 3.1 | 5.3 |
|  |  | Mother | 19 | 3.5 | 6.3 |

**Figure 3:**
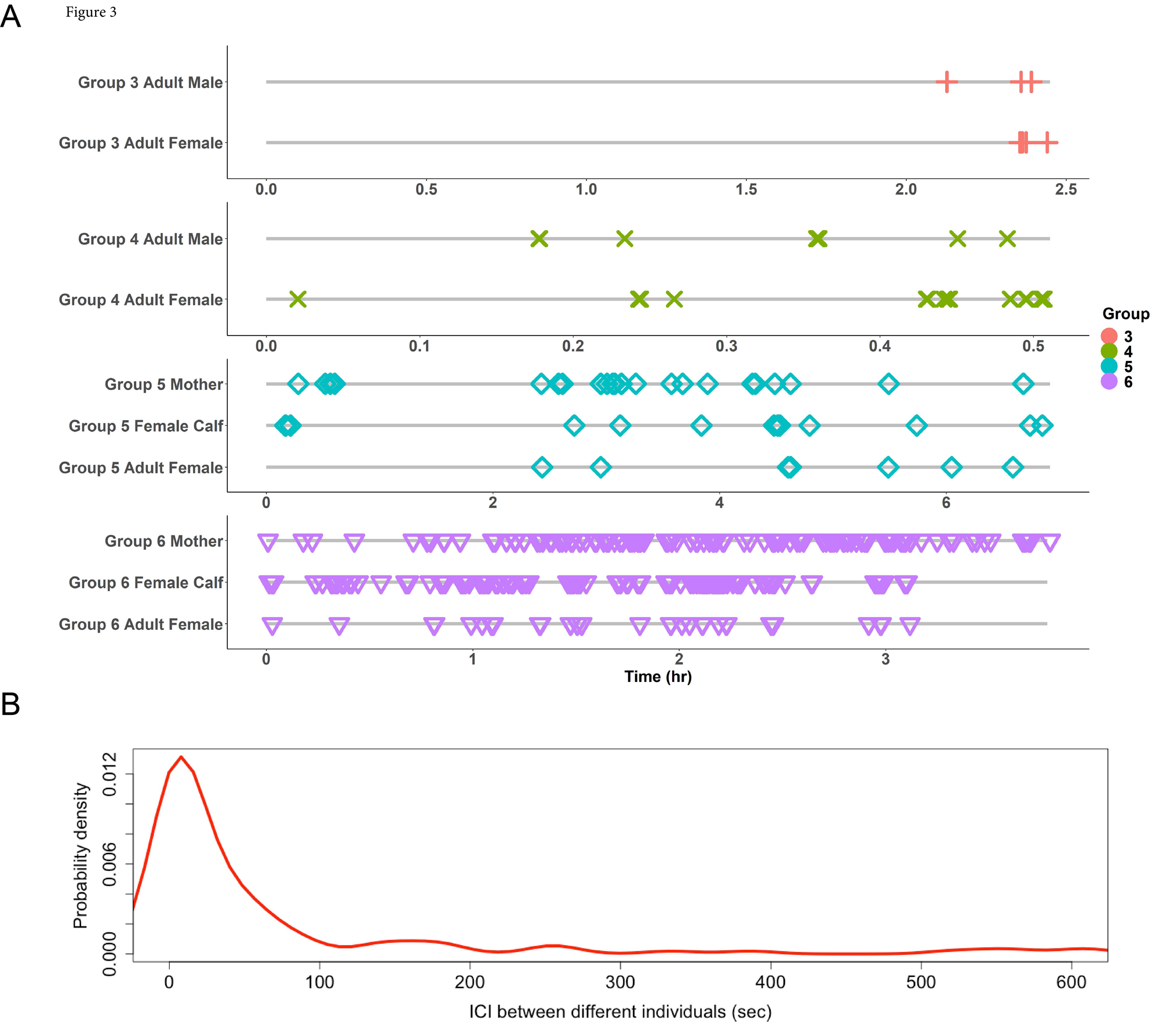
Relative timing of focal call production across individuals. A) Timelines of focal call occurrences (colored symbols) on each tag relative to the analysis period (gray line). Colors and symbols correspond to each different group. Groups 1, 2 and 7 are not shown because only one of the animals in the group vocalized. B) Probability density curve of the inter-call interval between different individuals. The area under the curve (AUC) from 0 to 100 seconds is 0.58

### Movement analysis

Tagged whales vocalized across the full range of dive depths observed on the tags (Figure 4). 13% of all calls were produced at/near the surface (i.e. less than 2m depth) and the rest were produced at various points during dives. The maximum depth of call production was 41 m, the minimum depth of call production was at the surface, and the mean depth of call production was 11 m (+/-7 m SD). Maximum dive depths ranged between 30 and 60 m and mean maximum dive depth across individuals was 45 m. The average water depth at the location of the tag deployments was approximately 62 m across tags (minimum water depth: 33 m, maximum: 125 m). There were no differences in depth of call production between calves and adults, although all groups with calves were tagged in water depths of 30-40 m, while adult-only groups were tagged in 60-125 m water depths.

**Figure 4:**
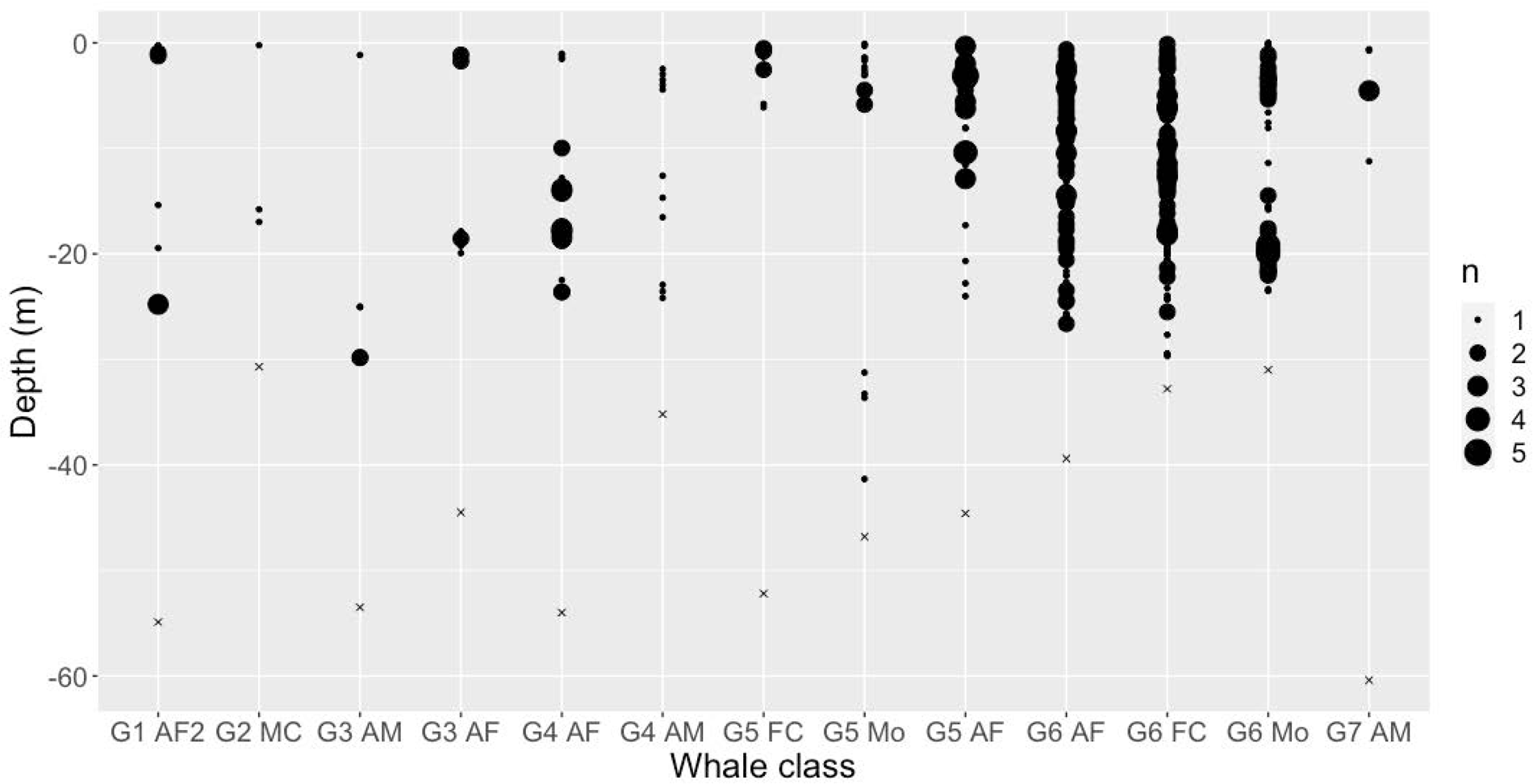
Depth of call production for all focal calls for each individual. Point size represents the number of calls at that depth and Xs mark maximum dive depth for that individual. Whale class is abbreviated to group (G) and number, plus two letters to mark sex, female reproductive status, and age class. M and F are used to denote male and female, A and C are used to denote adult and calf, and Mo denotes a mother. A number was added at the end when needed.

## Discussion

Using this approach of comparing relative received levels of calls recorded across tags on all whales in a group, we successfully assigned calls to callers for approximately 97% of calls in the dataset. Both focal and non-focal calls were recorded over a wide range of RLs, and the low end of the focal RL range was lower than that of the non-focal RL range. This indicates that although simultaneous tags often show a clear difference in call RLs across tags, and even though the distribution of non-focal RLs overlaps mostly with the lower end of the distribution of focal RLs, focal and non-focal calls occupy similar RL levels within a single tag. This is likely because whales vocalize at varying source levels both within and across call types, as has been described for humpback whale song (Stimpert et al. 2020). There still is a level of uncertainty with RL measurements, as there are differences in tag location which may impact tag differences in RL and other propagation effects may cause RL to vary depending on the environment. Thus, the range of RLs shown here is meant to be representative but could still reflect these measurement uncertainties. The range of RL results for focal and non-focal tags provide additional evidence that while an SNR threshold for determining focal calls may work in some cases, it may not always be robust enough to distinguish between focal and non-focal calls. This is likely true for other taxa as well since it is common for animal vocalizations to vary in amplitude across individuals and across contexts within an individual (Gustison and Townsend 2015).

After assigning focal calls based on relative received level, we were able to calculate both the call rate at both the individual and group levels. Since some calls were classified as indeterminate, actual call rates may be higher than our estimates, but this would primarily impact those tags with already high call rates. Call rate varied widely across individuals, with a mean individual call rate of about 23 calls per hour, but with some tag records that did not contain any vocalizations. Similarly, group-level call rate varied, with a mean group call rate of 55 calls per hour, but with some group call rates as low as 1 call per hour. Since humpback whales also seem to produce most of their calls in bouts (79% calls produced in bouts), call rate is not evenly distributed across recording time. It may be useful for future studies to report additional statistics such as bout rate and average bout length to better represent call rate over time. Call rate and bout rate have important implications for passive acoustic monitoring, and particularly for passive acoustic density estimation (Marques et al. 2013). We also found coordination in calling activity, meaning that most of the time, when one individual vocalizes, another individual responds. This behavior implies that call rate is likely dependent on social context, which is also important to consider in interpreting passive acoustic data, especially for density estimation.

Our bout production results are in alignment with previous results from this species on migration in Australia (Rekdahl et al. 2015, Cusano et al. 2022). Growing evidence of bout production by humpback whales across populations and habitats suggests that more research should investigate the social and behavioral context of these bouts. Additional data will allow for the development of functional hypotheses as well as an understanding of bout characteristics like syntax and rhythm and how these aspects of social call bouts compare to humpback whale song. In other species, acoustic sequences have been found to contain information related to signaler identity or context (e.g., Koren and Geffen 2012, Cäsar et al. 2013). Understanding the content and function of acoustic sequences is a growing area of research in animal behavior, and there is ongoing development of analytical techniques for answering questions related to acoustic sequences (Kershenbaum et al. 2016). We found a bout end criterion of 2.2 seconds, which, along with the previously calculated BEC of 3.9 seconds from the South Pacific (Rekdahl et al. 2015), means that humpback whales are producing bouts with short inter-call intervals. Inter-call intervals in vocal bouts may encode additional information, and in some cases may be indicative of social situations and arousal (Fischer et al. 1995, Handel et al. 2009). Humpback whale songs exhibit variable inter-unit intervals that on average range from about 0.5 to 2.5 s (Handel et al. 2009, Schneider and Mercado 2019). Thus, silent durations between sounds are similar in humpback whale social call bouts and song, although song inter-unit intervals may tend to be shorter. In contrast, the inter-unit intervals in blue whale songs are between about 5 and 14 seconds on average (Miller et al. 2014).

Since vocal exchanges are challenging to study without caller identification, this study is novel in our investigation of the timing of vocal production between individuals in this species. We found evidence that humpback whales are regularly calling back and forth with inter-individual call intervals of 100 seconds or less. Timing in vocal exchanges can indicate cooperative and turn-taking dynamics and mechanisms (Takahashi et al. 2013, Demartsev et al. 2018), or can encode information like dominance or internal state (Gamba et al. 2016, Fischer et al. 1995). In pygmy marmosets, the measured median time interval in vocal exchanges is about 5 seconds, which matches coupled oscillator dynamic predictions (Takahashi et al. 2013). Future research can investigate the dynamics of humpback whale vocal exchanges in more depth and test hypotheses related to information contained in call timing as well as the mechanisms underlying call timing, like coupled oscillator dynamics or other models as demonstrated in other taxa, including pygmy marmosets (Takahashi et al. 2013), meerkats (*Suricata suricatta,* Demartsev et al. 2018), and humans.

An additional factor that is important for passive acoustic monitoring is the depth at which marine animals are calling. We found that humpback whales are calling at various depths throughout their dives in this shallow habitat. In contrast, right whales predominantly signal near the surface (Parks et al. 2011) and blue whales have been found to predominantly call at shallow depths (<30m), even while making deep dives (>100m, Oleson et al. 2007). Short-finned pilot whales vocalize both while socializing at the surface and during deep (up to 800m) foraging dives (Jensen et al. 2011). For humpback whales, evidence of call production throughout the water column may indicate the use of vocalizations across different behavioral contexts (i.e., coordinated foraging, social interaction) across depths, and future research could further investigate behavioral context and function of different call types relative to location in the water column. This is useful for understanding risk for anthropogenic disturbance like entanglement or ship strikes, as well as for modeling acoustic propagation and detection range of vocalizations for acoustic monitoring.

Simultaneously equipping all the individuals in a social group with recorders has the potential to be useful across taxa for studies of individual and group-level acoustic behavior and facilitates the study of social interactions. Where possible, future studies requiring robust caller identification can prioritize deploying tags on all the animals in a group to compare call received level across recorders. However, future research on additional methods for differentiating individual callers in acoustic data remains important. Deploying acoustic recorders on all individuals in a group can be restrictive, especially when social context changes frequently or when group size exceeds the number of tags available for deployment. An additional requirement of simultaneous tagging for caller ID is concurrent behavioral observations to track social affiliations.

This study provides evidence of the feasibility of using simultaneous tag data for caller identification with small groups of baleen whales and offering a more robust method for identifying focal calls than an SNR threshold. It will also be useful for future studies to pair this simultaneous tag method with analysis of accelerometer records for signatures of vocalizations (as in Goldbogen et al. 2014, Saddler et al. 2017, Stimpert et al. 2020) and thus cross-validate different methods for identifying calls from tagged baleen whales. Using this method, we were able to gain insight into individual humpback whale acoustic behavior, including a description of inter-call intervals between and within individuals, which provides preliminary baseline data that can be used for future research related to rhythm, sequence production, and cooperative behavior. These data also allowed for the calculation of call rates and call production as it relates to dive behavior, which will be useful for conservation applications including passive acoustic monitoring and density estimation.

## Acknowledgments

Tag data were collected under National Marine Fisheries Service permits 775-185 (to Northeast Fisheries Science Center) and 605-1904 (to Whale Center of New England). We would like to thank all of the members of the field team from 2006-2009 and the crews of the NOAA R/Vs Nancy Foster and Auk, and the many individuals involved in the long-term cataloging of humpback whales in the Gulf of Maine.

## Author contributions

Conceptualization: JMZ, SEP, FHJ, VPM; Methodology: FHJ, JMZ, VPM, Software: FHJ, JMZ; Validation: JMZ, VPM, DLA; Formal analysis: JMZ, VPM, DLA, KJK; Investigation: DNW, SEP, JR, JET, MW, ASF; Resources: DNW, SEP, JR, ASF; Data curation: JMZ, VPM, DLA, KJK, JR, MW; Writing – original draft preparation: JMZ; Writing – review and editing: JMZ, VPM, DLA, FHJ, KJK, JR, JET, ASF, MW, DNW, SEP; Visualization: JMZ, DLA; Supervision: SEP; Project administration: JMZ; Funding acquisition: SEP, FHJ, DNW, JR, MW

## Data availability

Data are available upon request.

